# 5β-Dihydrotestosterone reveals a mutant androgen receptor vulnerability in prostate cancer

**DOI:** 10.64898/2026.07.14.738538

**Authors:** Samual Adams, Lindsey Phelan, Therese Lewis, Josie Behm, Aggie Law, Xianglin Shi, Guangyuan (Frank) Li, Jianneng Li

## Abstract

Bipolar androgen therapy (BAT) exploits the paradoxical vulnerability of castration-resistant prostate cancer (CRPC) cells to rapid cycling between castrate and supraphysiologic androgen concentrations, but clinical BAT uses testosterone, which can also activate wild-type androgen receptor (AR) in androgen-responsive tissues, causing systemic side effects. 5β-dihydrotestosterone (5β-DHT) is a naturally occurring testosterone metabolite generally considered androgenically inactive because it binds wild-type AR weakly, yet its activity against clinically relevant AR mutants has not been systematically evaluated. Here, we tested whether 5β-DHT and related 5β-reduced testosterone metabolites activate AR signaling and growth programs in prostate cancer models that carry AR mutations. In C4-2 cells, 5β-DHT and 3β-etiocholanediol (3β-ecdiol) increased canonical AR target genes, including *KLK3* and *TMPRSS2*, with weaker activity than testosterone, whereas other 5β metabolites showed limited activity. In androgen-responsive LNCaP and C4-2 models, 5β-DHT and 3β-ecdiol promoted cell growth under androgen-depleted conditions, and this effect was suppressed by enzalutamide, supporting AR dependence. RNA-seq confirmed that 5β-DHT and 3β-ecdiol induced androgen-response gene sets substantially overlapping with testosterone, albeit at lower transcriptional magnitude. Further, we found that 5β-DHT, but not 3β-ecdiol, suppresses cell proliferation of LNCaP, C4-2, and PC-3 cells stably expressing the clinically relevant AR gain-of-function mutants W742C and H875Y through activating AR-induced senescence-like features after high-dose exposure, consistent with the therapeutic logic of BAT. These findings identify 5β-DHT as an overlooked mutant-AR agonist capable of BAT-like tumor suppression and propose it as a testosterone surrogate in BAT with potentially reduced systemic androgenic side effects.

**Highlights:**

- 5β-DHT and 3β-ecdiol promote AR-dependent prostate cancer cell growth
- Both are weaker AR agonists than testosterone by RNA-seq and qPCR
- Supraphysiologic 5β-DHT suppresses growth via AR-mediated senescence
- Growth suppression extends to AR mutants W742C and H875Y
- 5β-DHT may be a lower-androgenicity testosterone surrogate for BAT

## 1. Introduction

Androgen receptor (AR) signaling is central to prostate cancer biology. The clinical foundation of androgen-deprivation therapy derives from the observation that metastatic prostate cancer regresses when androgen action is suppressed and can worsen when androgen is restored [1]. Despite initial responses, most advanced tumors eventually progress to castration-resistant prostate cancer (CRPC), a state that frequently remains dependent on AR signaling through mechanisms that include AR amplification, intratumoral androgen synthesis, ligand-independent activation, splice variants, and ligand-binding-domain mutations [2–7]. The persistence of AR signaling in CRPC has enabled next-generation AR pathway antagonists, such as abiraterone and enzalutamide, which extend survival, but resistance occurs through overlapping AR-driven mechanisms and AR bypass [8–14], leaving patients with limited options.

Paradoxically, supraphysiological androgen can suppress CRPC through AR-dependent mechanisms that underlie bipolar androgen therapy (BAT) [15–17]. Chronic androgen deprivation selects tumors with elevated AR expression or activity, making them vulnerable to acute androgen excess [15, 16]. High-dose androgen can disrupt DNA replication licensing, induce AR-dependent DNA damage, suppress MYC- and cell cycle-associated signaling, and then trigger cell-cycle arrest, apoptosis, or senescence [16–28]. BAT exploits these effects by rapidly cycling androgen levels between supraphysiological and castrate states, thereby stressing AR-addicted tumor cells while limiting adaptive escape. Clinical trials have shown that BAT has meaningful activity in subsets of men with metastatic CRPC and can restore sensitivity to subsequent AR-directed therapy [17, 29–32]. However, BAT relies on testosterone (T), a broadly active androgen that can stimulate wild-type AR in nonmalignant androgen-responsive tissues, leading to polycythemia, cardiovascular risk, and systemic androgenic effects [33]. Therefore, a T substitute that preserves the ability to stress mutant or overexpressed AR in CRPC while reducing wild-type AR activation could improve the therapeutic window of BAT.

Testosterone metabolism generates two major 5-reduced steroids with distinct biological properties. Conversion to 5α-dihydrotestosterone (5α-DHT) produces a potent androgen with high AR affinity [34, 35], whereas 5β-dihydrotestosterone (5β-DHT) has historically been considered non-androgenic or genomically inactive because its cis A/B-ring junction creates a bent steroid conformation that alters ligand geometry and has been regarded as poorly compatible with canonical wild-type AR ligand-binding domain occupation [36, 37]. A critical unresolved question is whether 5β-DHT is truly inactive across the altered AR landscape of CRPC. Ligand-binding domain mutations such as T878A, H875Y, and W742C can broaden steroid recognition and alter responses to antiandrogens or noncanonical ligands [38, 39]. Because these mutations reshape ligand specificity, a steroid that is weak at wild-type AR may still activate mutant AR sufficiently to affect prostate cancer biology [38, 39].

In the present study, we investigated 5β-DHT and related testosterone metabolites in AR-responsive prostate cancer models harboring AR mutations and in engineered mutant-AR models. We find that 5β-DHT is not inert: it activates AR-regulated transcription, induces growth in AR-dependent prostate cancer cells by engaging androgen-response programs, and produces high-dose senescence-like phenotypes, while activating AR less potently than T. These findings provide a mechanistic rationale for evaluating 5β-DHT as a potential lower-androgenicity substitute for testosterone in BAT with potentially reduced systemic androgenic side effects.

## 2. Materials and Methods

### Cell lines, plasmids, and chemicals

C4-2 and LNCaP (AR-positive, androgen-sensitive), and PC-3 (AR-null) human prostate cancer cells (ATCC) were maintained in RPMI-1640 (Cytiva) supplemented with 10% or 5% FBS (GeminiBio) and 1% penicillin-streptomycin (Corning). Clinically relevant AR mutants (W742C and H875Y) were generated using the Q5 Site-Directed Mutagenesis Kit (New England Biolabs) with the pLENTI6.3/AR-GC-E2325 (a gift from Karl-Henning Kalland, Addgene plasmid # 85128 [40]) template. Primers were designed via NEBbasechanger: W742C: F-TCAGTACTCCTGCATGGGGCTCA, R-ATGACAGCCATCTGGTCG; H875Y: F-GAGAGAGCTGTATCAGTTCACTTTTG, R-GCAATAGGCTGCACG. Mutant and WT AR Lentiviral particles were produced in 293T cells using the AR plasmids, pMD2.G and psPAX2 vectors with the Hieff Trans™ Liposomal Transfection Reagent (Yeason) and then transduced into PC3 cells as previously described [12, 13]. Stable cell lines were selected using Blasticidin (3 µg/mL). For low-androgen experiments, cells were transferred to medium supplemented with 10% charcoal-stripped FBS (csFBS, GeminiBio) for 48 h before treatment. Full-serum (FBS) conditions were used to model the BAT context. All cells were maintained at 37°C in 5% CO_2_. Cell lines were authenticated by DDC Medical and routinely screened for mycoplasma contamination using primers 5’-ACACCATGGGAGCTGGTAAT-3’ and 5’-GTTCATCGACTTTCAGACCCAAGGCAT-3’ [12, 13]. Testosterone (T), 5β-DHT, 3β-ecdiol, 3β-ecl, 3α-ecdiol, 3α-ecl, 5β-ecdione, pregnenolone, progesterone, 5α-dihydroprogesterone, allopregnanolone, and their 17α-hydroxy derivatives were obtained from Steraloids and dissolved in ethanol. Enzalutamide (Enz, MedChemExpress) was used at 10 μM as a pharmacological AR antagonist control.

### Cell proliferation assays

C4-2, LNCaP, PC-3 parental cells, and PC-3 cells stably expressing AR-W742C or AR-H875Y (2,000/well) were seeded onto poly-L-ornithine (PLO, Sigma-Aldrich) coated 96-well plates in media containing FBS or csFBS. Following overnight attachment, cells were treated in triplicate with the indicated conditions. Longitudinal imaging was performed every 8 hours for 10 days using a CellCyteX (ECHO/Cytena) live cell imager. Percent confluence was used as a proxy for cell number and expressed as a fold change relative to time zero.

### Gene expression and immunoblot

Total RNA was extracted using the Total RNA extraction kit (ELK Biotechnology) and reverse transcribed using Hifair™ III 1st Strand cDNA Synthesis Super Mix (Yeason). Quantitative PCR was performed on QuantStudio 3 (Thermo Fisher Scientific) in triplicate using the following primers: *RPLP0*: F-TGGCAATCCCTGACGCAC, R-CACGTTGTCTGCTCCCACAAT; *KLK3*: F-AGAGCTGTGTCACCATGTG, R-AGGGTTGGGAATGCTTCTC; *TMPRSS2*: F-GGAGTGTACGGGAATGTGATG, R-CCAGCCCCATTGTTTTCTTG; *ALDH1A3*: F-CTAACGGGGCCGTGGAAAA, R-TCCCACTCTTGGATTCGTGC; *RRAS*: F-CGGCGTGGGCAAGAGC, R-ACTGCAGATCTTCGTGTAGGAGT; *CDKN1A*: F-TTGTACCCTTGTGCCTCGCT, R-GTGGTAGAAATCTGTCATGCTGG; *CDKN2B*: F-CGCGGGGACTAGTGGAGAAG, R-CCCATCATCATGACCTGGATCG; *LMNB1*: F-TCTTGCTACTGCACTTGGTGA, R-AAGGCTCTGACAACGATTCTCC; *CCNA2*: F-TGGACCTTCACCAGACCTACC, R-GTGGGTTGAGGAGAGAAACAC. Each mRNA transcript was quantified by normalizing the sample values to *RPLP0* and to vehicle-treated cells. All gene expression studies were repeated in at least three independent experiments.

For protein analysis, immunoblots were performed as described previously [12, 13]. Briefly, cells were seeded onto PLO-coated 12-well plates in 10% FBS RPMI and treated with EtOH (vehicle), 100 nM Testosterone, or 5β-DHT for 1 or 3 days. Cells were harvested in SDS lysis buffer. Proteins were resolved by SDS-PAGE and transferred to PVDF membranes for immunoblotting. After incubating with the anti-p-RB1 (ser807/811) (Proteintech-30376-1-AP), anti-p57 (Proteintech-66794-1-Ig), anti-SKP2 (Proteintech-15010-1-AP), anti-p27 (Proteintech-25614-1-AP), anti-p21 (Proteintech-10355-1-AP), and β-actin (Engibody-AT0001) overnight at 4°C, the appropriate secondary antibody was incubated for 1 hour at room temperature. A chemiluminescent detection system (Engibody) was used to detect bands with peroxidase activity, and a Bio-Rad Chemidoc was used for imaging.

### RNA-sequencing

C4-2 cells were treated with 10 nM T, 5β-DHT, or 3β-ecdiol in csFBS medium for 24 h, or with 100 nM T or 5β-DHT in FBS medium for Day 1 (D1) or Day 3 (D3). RNA was extracted using the ELK Total RNA Extraction Kit or Zymo Quick-RNA Miniprep kit. Purified RNA (normalized to 50 ng/µL) was submitted to Plasmidsaurus (Eugene, OR) for library preparation and sequencing. Briefly, mRNA was converted to cDNA by reverse transcription using a poly(dT)VN primer, followed by second-strand synthesis, tagmentation, library indexing with unique dual indices (UDIs), and amplification. Libraries were sequenced on an Illumina platform using a single-end, stranded 3’ end counting approach (∼90 bp reads) with unique molecular identifiers (UMIs) for PCR deduplication, targeting ≥10 million deduplicated reads per sample. Raw sequencing reads were demultiplexed using BCL Convert v4.3.6 and fqtk v0.3.1, then quality-filtered with FastP v0.24.0. Filtered reads were aligned to the human reference genome GRCh38 using STAR v2.7, coordinate-sorted with samtools v1.21, and UMI-deduplicated using UMICollapse v1.1.0. Gene-level counts were generated using featureCounts from subread v2.1.1. Differential expression analysis was performed for each treatment relative to its matched control using edgePython v0.2.5 with TMM normalization. Significantly regulated genes were defined using adjusted p < 0.05, |log FC| ≥ 1. Gene set enrichment analysis was performed using GSEApy v0.12 on pre-ranked gene lists (ranked by log2 fold-change) using the MSigDB Hallmark and selected pathway gene sets; this pre-ranked approach is compatible with 3’ end counting data, in which gene-level rankings reflect transcript abundance estimates rather than full-transcript read coverage. Heatmaps were generated from normalized expression values scaled by row z-score. RNA-seq data will be deposited in NCBI GEO upon acceptance.

### GFP-ARE reporter assay

PC3 cells were first transduced with the ARR3tk-eGFP/SV40-mCherry reporter (a gift from Charles Sawyers, Addgene plasmid # 132360 [41]). RFP-positive cells were isolated via FACS in the Harper Cancer Research Institute core facility and single-colony selection before being used to generate AR W742C or AR H875Y stable cell lines. Cells were seeded in medium with 5% csFBS, cultured for 48 hours, and then treated with the indicated conditions. GFP and RFP expressions were monitored for 48 hours via the green and red channels of a CellCyteX imager. AR activation was quantified longitudinally as the GFP+/RFP+ cell ratio.

### SA-β-gal staining

Senescence-associated β-galactosidase activity was detected with the Cell Senescence Staining Kit (Cell Signaling Technology) at pH 6.0 per the manufacturer’s protocol. Images were acquired using a Leica Matero FL and analyzed with ImageJ.

### Statistics

Data are presented as mean ± SEM from at least three independent experiments. Comparisons were performed using one-way ANOVA with Tukey’s post-hoc test or Student’s two-tailed t-test as appropriate. *p < 0.05, **p < 0.01, ***p < 0.001, ****p < 0.0001.

## 3. Results

### 3.1. 5β-DHT and 3β-ecdiol promote AR-dependent C4-2 and LNCaP cell growth through AR activation

Testosterone can be metabolized by 5β-reductase (SRD5B1) in the liver, which generates 5β-DHT. Aldo-keto reductase 1C (AKR1C) and 17β-hydroxysteroid dehydrogenases (HSD17B) enzymes further convert 5β-DHT to a series of etiocholanolone metabolites: 3β-etiocholanediol (3β-ecdiol), 3β-etiocholanolone (3β-ecl), 3α-etiocholanediol (3α-ecdiol), 3α-etiocholanolone (3α-ecl), and 5β-etiocholanedione (5β-ecdione) (Fig. 1A).

**Fig. 1.**
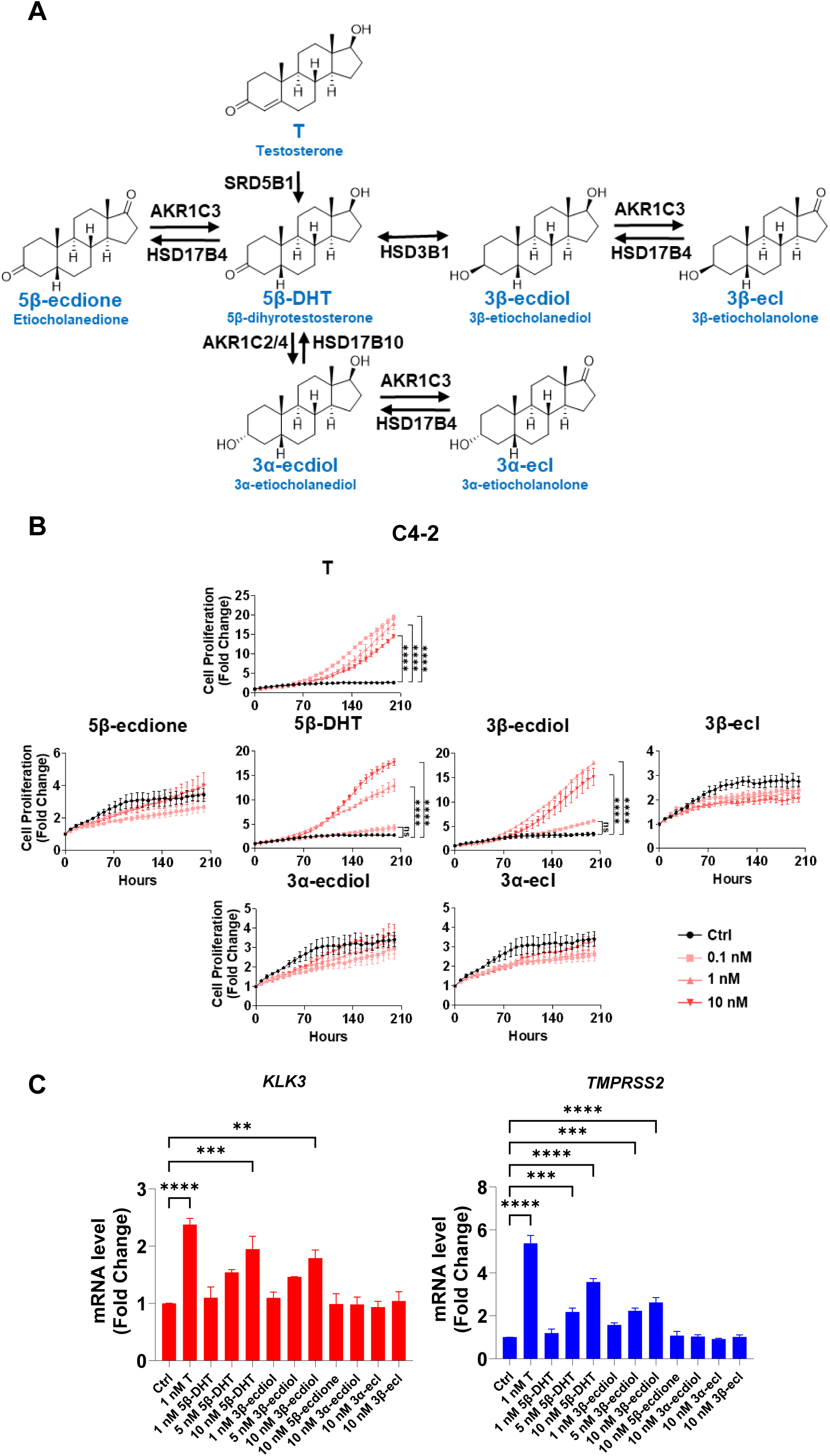
5β-DHT and 3β-ecdiol promote AR-dependent C4-2 cell growth and induce AR target genes in androgen-depleted conditions. (**A**) Schematic of the T metabolic pathway to 5β-DHT (via SRD5B1) and downstream etiocholanolone metabolites via AKR1C2/4, AKR1C3, HSD3B1, and HSD17B4. (**B**) Cell proliferation curves of C4-2 cells in csFBS medium treated with T, 5β-ecdione, 5β-DHT, 3β-ecdiol, 3β-ecl, 3α-ecdiol, or 3α-ecl (0.1, 1, 10 nM) for up to 210 h; fold change relative to time zero. (**C**) RT-qPCR of *KLK3* and *TMPRSS2* mRNA (5β-DHT: 1, 5, 10 nM; T: 1 nM; 3β-ecdiol: 1, 10 nM; others: 10 nM). Data are mean ± SEM (n ≥ 3). **p < 0.01, ***p < 0.001, ****p < 0.0001 vs. Ctrl.

To determine whether these testosterone metabolites exhibit androgenic activity in AR-mutant cells, we treated C4-2 cells, which harbor the AR T878A mutation, with T or each of the six 5β-metabolites at 0.1, 1, and 10 nM in csFBS medium and monitored proliferation by live-cell imaging (Fig. 1B). T promoted concentration-dependent C4-2 cell growth as expected. Among the six metabolites, only 5β-DHT and 3β-ecdiol significantly stimulated proliferation above vehicle control at all three concentrations; 5β-ecdione, 3β-ecl, 3α-ecdiol, and 3α-ecl had no effect at any dose tested (Fig. 1B). The growth-promoting activity of 5β-DHT was intermediate between T and 3β-ecdiol, establishing a potency ordering consistent with progressive metabolic inactivation. RT-qPCR analysis confirmed that 5β-DHT and 3β-ecdiol upregulated the canonical AR target genes *KLK3* and *TMPRSS2* in C4-2 cells in a concentration-dependent manner, but at mRNA levels below those induced by equimolar T, indicating authentic but attenuated AR transcriptional activation (Fig. 1C).

To verify the generalizability and AR dependency of these findings, we tested the same compounds in LNCaP cells and in AR-null PC-3 parental cells. Both 5β-DHT and 3β-ecdiol promoted LNCaP proliferation in csFBS medium, mirroring the C4-2 results (Fig. S1A). In contrast, neither compound stimulated growth of AR-null PC-3 cells, confirming strict AR dependency (Fig. S1B). Pharmacological validation was provided by co-treatment with ENZ (10 μM), which fully abrogated the growth-promoting effects of both 5β-DHT and 3β-ecdiol in C4-2 cells (Fig. S1C). Together, these data show that 5β-DHT and 3β-ecdiol are not merely inactive terminal metabolites of testosterone: in an androgen-depleted context, they can function as weak AR agonists capable of sustaining growth of AR-dependent prostate cancer cells.

### 3.2. Genome-wide transcriptomics reveal weaker but authentic androgen response activation by 5β-DHT and 3β-ecdiol

To define the genome-wide transcriptional consequences of 5β-DHT and 3β-ecdiol exposure, we performed RNA sequencing of C4-2 cells treated with 10 nM testosterone, 5β-DHT, or 3β-ecdiol for 24 hours in csFBS medium. Gene set enrichment analysis (GSEA) using the MSigDB Hallmark gene set collection revealed that the Androgen Response pathway was the top-enriched gene set across all three treatments relative to vehicle control, confirming that T, 5β-DHT, and 3β-ecdiol each activate canonical AR-driven transcription (Fig. 2A). Beyond the androgen response, 5β-DHT and 3β-ecdiol co-enriched a broader set of proliferative pathways, including G2M Checkpoint, MYC Targets V1/V2, MTORC1 Signaling, Mitotic Spindle, and E2F Targets, consistent with their growth-promoting activity in androgen-depleted conditions. Differential gene expression analysis (DEGs: adjusted p < 0.05, absolute log2 fold change ≥ 1) revealed a clear hierarchy: T induced the broadest response (768 DEGs), 5β-DHT activated a substantially smaller set (190 DEGs, of which 174 overlapped with T), and 3β-ecdiol induced only 36 DEGs (all shared with T) (Fig. 2B). The small number of 5β-DHT-unique DEGs (16 genes) and complete overlap of 3β-ecdiol-regulated genes with T-regulated genes suggest that these compounds do not activate novel transcriptional programs but rather engage the same AR-driven regulon as T at reduced amplitude.

**Fig. 2.**
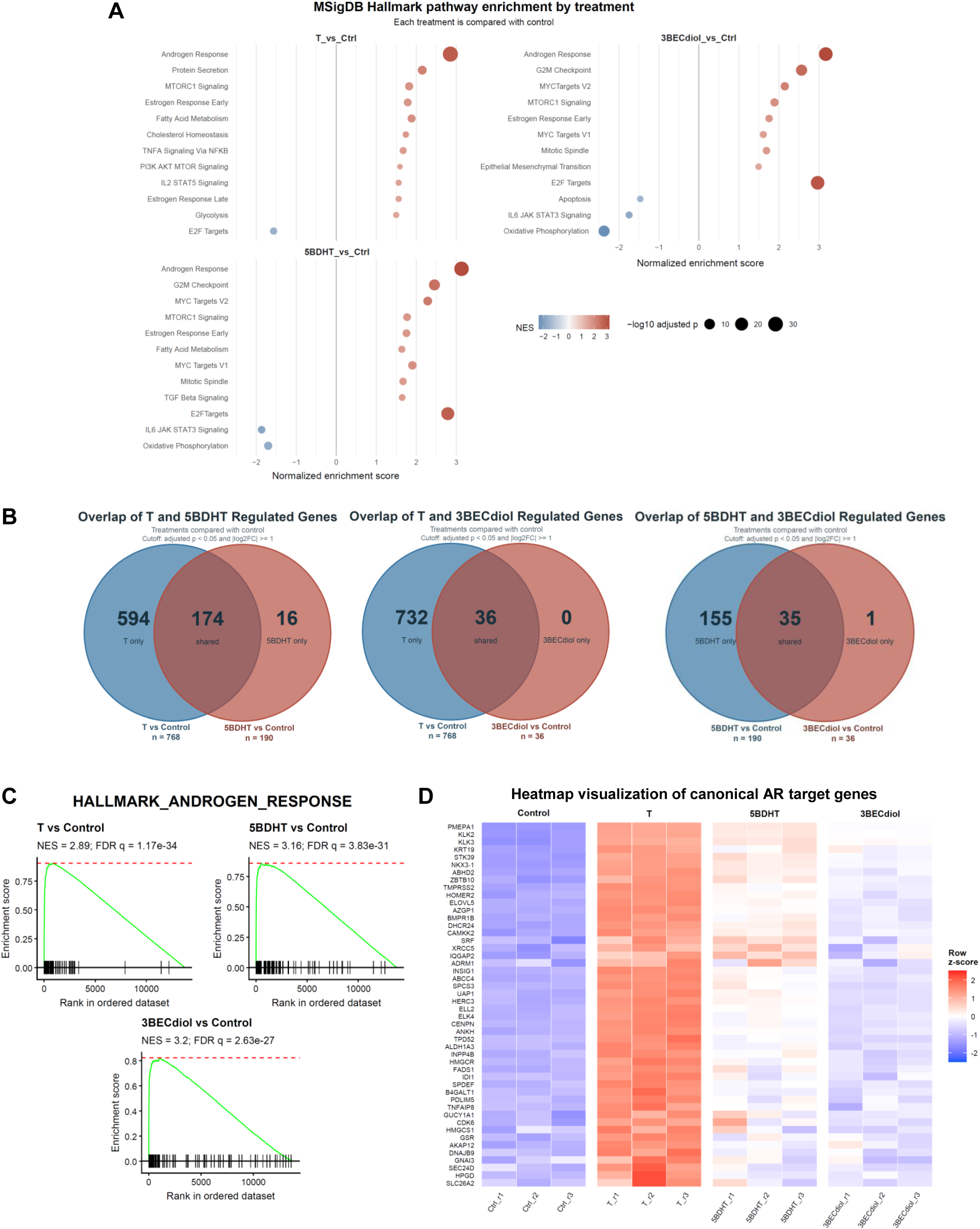
Genome-wide transcriptomics confirms androgen response activation by 5β-DHT and 3β-ecdiol with a smaller DEG footprint than T. C4-2 cells in csFBS medium were treated with 10 nM T, 5β-DHT, or 3β-ecdiol for 24 h (n = 3 per condition). (**A**) MSigDB Hallmark GSEA dot plots for each treatment vs. vehicle. Dot size: −log10 adjusted p; color: NES. (**B**) Venn diagrams of DEGs (adjusted p < 0.05, |log2FC| ≥ 1): T (768), 5β-DHT (190), 3β-ecdiol (36). (**C**) GSEA enrichment plots for HALLMARK_ANDROGEN_RESPONSE. (**D**) Heatmap of canonical AR target genes (row z-score).

Moreover, GSEA enrichment plots for the HALLMARK_ANDROGEN_RESPONSE gene set confirmed robust and statistically significant enrichment by all three treatments relative to vehicle control, with comparably high normalized enrichment scores: T (NES = 2.89, FDR q = 1.17×10□³□), 5β-DHT (NES = 3.16, FDR q = 3.83×10 ³¹), and 3β-ecdiol (NES = 3.2, FDR q = 2.63×10□²□) (Fig. 2C). The steep leading-edge enrichment pattern observed in all three plots indicates that androgen response genes are concentrated at the top of each ranked gene list, consistent with direct AR transcriptional activation by 5β-DHT and 3β-ecdiol at a level qualitatively comparable to T (Fig. 2C). Heatmap visualization of canonical AR target genes, including *FKBP5*, *KLK2*, *KLK3*, *STEAP4*, *NKX3-1*, and *TMPRSS2*, confirmed directionally concordant but quantitatively attenuated responses to 5β-DHT and 3β-ecdiol relative to T (Fig. 2D). Collectively, these results position 5β-DHT and 3β-ecdiol as an attenuated testosterone mimetic at the AR transcriptional level.

### 3.3. Supraphysiologic 5β-DHT suppresses CRPC growth including AR gain-of-function mutant lines

Given that both 5β-DHT and 3β-ecdiol activate mutant AR at nanomolar concentrations, we hypothesized that supraphysiologic doses would suppress growth via a BAT-like mechanism. C4-2 and LNCaP cells in FBS medium were treated with T, 5β-DHT, or 3β-ecdiol at 100 nM or 1 μM (Fig. 3). In both AR mutant prostate cancer cell lines, both 100 nM and 1 μM T markedly suppressed proliferation relative to vehicle control, recapitulating the BAT growth-suppressive effect. Critically, high-dose 5β-DHT, but not 3β-ecdiol, significantly suppressed C4-2 and LNCaP cell growth in a dose-dependent manner (Fig. 3A and 3B); given that 5β-DHT is a more potent AR agonist than 3β-ecdiol, as evidenced by its greater number of differentially expressed genes and broader hallmark pathway enrichment at equimolar concentrations (Fig. 2), this differential growth-suppressive activity suggests that a minimum threshold of AR agonist potency is required to trigger the BAT-like growth-suppressive effect, and that 5β-DHT, but not the weaker agonist 3β-ecdiol, meets this threshold, supporting its potential as a testosterone substitute for BAT.

**Fig. 3.**
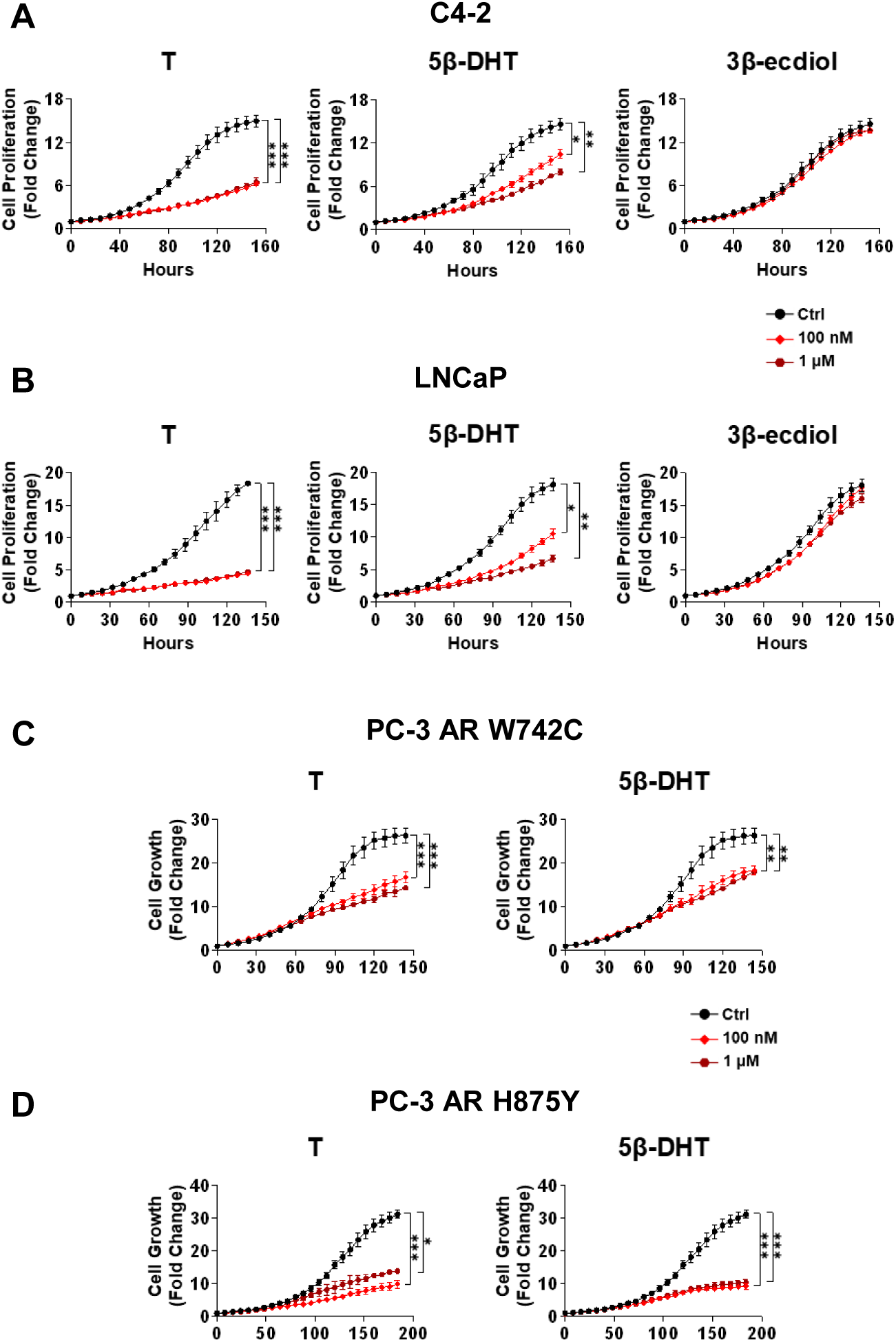
Supraphysiologic 5β-DHT suppresses CRPC growth in C4-2, LNCaP, and AR mutant overexpressing PC-3 cells. Proliferation of C4-2 (**A**) and LNCaP (**B**) in FBS medium with T, 5β-DHT, or 3β-ecdiol at Ctrl, 100 nM, or 1 μM. (**C**) Growth of PC-3 AR W742C and PC-3 AR H875Y cells treated with T or 5β-DHT at Ctrl, 100 nM, or 1 μM in FBS medium. Data are mean ± SEM (n ≥ 3). *p < 0.05, **p < 0.01, ***p < 0.001.

We next assessed whether this BAT-like effect extended to cells expressing other clinically prevalent AR gain-of-function mutations, including H875Y and W742C. In PC-3 cells stably expressing AR W742C or AR H875Y, high-dose 5β-DHT suppressed cell growth comparable to T (Fig. 3C and 3D), supporting extension of the BAT-like effect to multiple mutant-AR contexts. To confirm that this growth suppression was accompanied by direct AR engagement, a dual GFP/RFP AR reporter system was stably introduced into PC-3 cells using the ARR3tk-eGFP construct [41], in which eGFP expression is driven by a probasin promoter modified to contain three androgen response elements [42], followed by stable overexpression of AR mutations. In AR W742C cells, both T and 5β-DHT increased the GFP+/RFP+ cell ratio in a time- and dose-dependent manner, with 1 μM producing greater activation than 100 nM for both compounds, though T was a stronger activator than 5β-DHT at each dose (Fig. S2A and S2C). In AR H875Y cells, testosterone and 5β-DHT both activated the AR-responsive reporter over time. The 100 nM and 1 μM doses produced largely similar responses for each ligand, while testosterone showed slightly stronger activation than 5β-DHT at 100 nM (Fig. S2B and S2D). Importantly, vehicle-treated cells showed no reporter activation in either mutant line, confirming ligand-dependent AR engagement by 5β-DHT across two clinically distinct gain-of-function AR mutations. Moreover, AR-null PC-3 parental cells were completely unaffected by supraphysiologic 5β-DHT and all other metabolites, confirming AR-dependent growth suppression (Fig. S3C). The four inactive metabolites (5β-ecdione, 3β-ecl, 3α-ecdiol, 3α-ecl) also failed to suppress C4-2 and LNCaP growth at 100 nM and 1 μM (Fig. S3A and S3B), establishing that AR agonist activity is required for the BAT effect.

To define whether the BAT-like effect is shared by other steroids that activate AR, we evaluated progesterone and seven structurally related progestins in C4-2 cells (Fig. S4A). Several compounds (pregnenolone, progesterone, 5α-DHP, 17OHP, and their 17α-hydroxy derivatives) promoted C4-2 proliferation in csFBS medium at low nanomolar doses, consistent with weak AR activation [39, 43] (Fig. S4B). However, none of these steroids significantly suppressed C4-2 growth at 100 nM and 1 μM in FBS medium (Fig. S4C). Despite promoting proliferation at physiologic doses, progesterone-class steroids failed to suppress growth at supraphysiological doses, distinguishing them from 5β-DHT. This difference suggests that progesterone and its analogs activate AR less potently than 5β-DHT and fail to reach the AR activation threshold required to trigger BAT-like growth suppression. This interpretation is further supported by the potency-threshold model: high-dose 5β-DHT, but not the weaker AR activator 3β-ecdiol, significantly suppressed cell growth. Together, these data demonstrate that the capacity to induce BAT-like growth suppression is not a general property of AR-activating steroids but rather depends on the magnitude of AR agonist potency, and that 5β-DHT uniquely meets this threshold among the testosterone metabolites and progesterone-class steroids examined here.

### 3.4. 5β-DHT recapitulates the androgen response and cellular senescence transcriptomes under BAT conditions

To characterize the transcriptional response to supraphysiologic 5β-DHT in the BAT context, we performed RNA-seq in C4-2 cells treated with 100 nM T or 5β-DHT in FBS medium at Day 1 and Day 3 (Fig. 4). GSEA confirmed that Androgen Response was the top Hallmark pathway at both time points for both compounds, with comparable NES values: T D1 (NES = 3.14, FDR q = 3.95×10□³□), 5β-DHT D1 (NES = 3.24, FDR q = 3.16×10□³□), T D3 (NES = 3.01, FDR q = 1.04×10□³¹), and 5β-DHT D3 (NES = 3.1, FDR q = 3.4×10□³²) (Fig. 4A and 4C). Concurrently, proliferative gene sets, including G2M Checkpoint, MYC Targets V1, and E2F Targets, were significantly negatively enriched (blue dots, NES < −2) under both T and 5β-DHT treatment at both time points, indicating suppression of cell cycle progression programs consistent with growth arrest (Fig. 4A). Additional pathways including Protein Secretion, TNFA Signaling Via NFKB, Fatty Acid Metabolism, and Hypoxia were positively enriched by both compounds, with the breadth and magnitude of enrichment increasing from Day 1 to Day 3, reflecting a progressive transcriptional reprogramming that parallels the deepening growth suppression observed over time (Fig. 4A). DEG analysis further revealed progressive transcriptional convergence between T and 5β-DHT: at D1, T induced 341 DEGs and 5β-DHT induced 85 DEGs with 80 shared; by D3, T induced 566 DEGs and 5β-DHT induced 241 DEGs with 213 shared, with heatmap analysis of leading-edge AR target genes confirming qualitatively identical but quantitatively attenuated responses to 5β-DHT compared with T at both time points, consistent with a deepening, AR-driven transcriptional commitment over time (Fig. 4B and 4D).

**Fig. 4.**
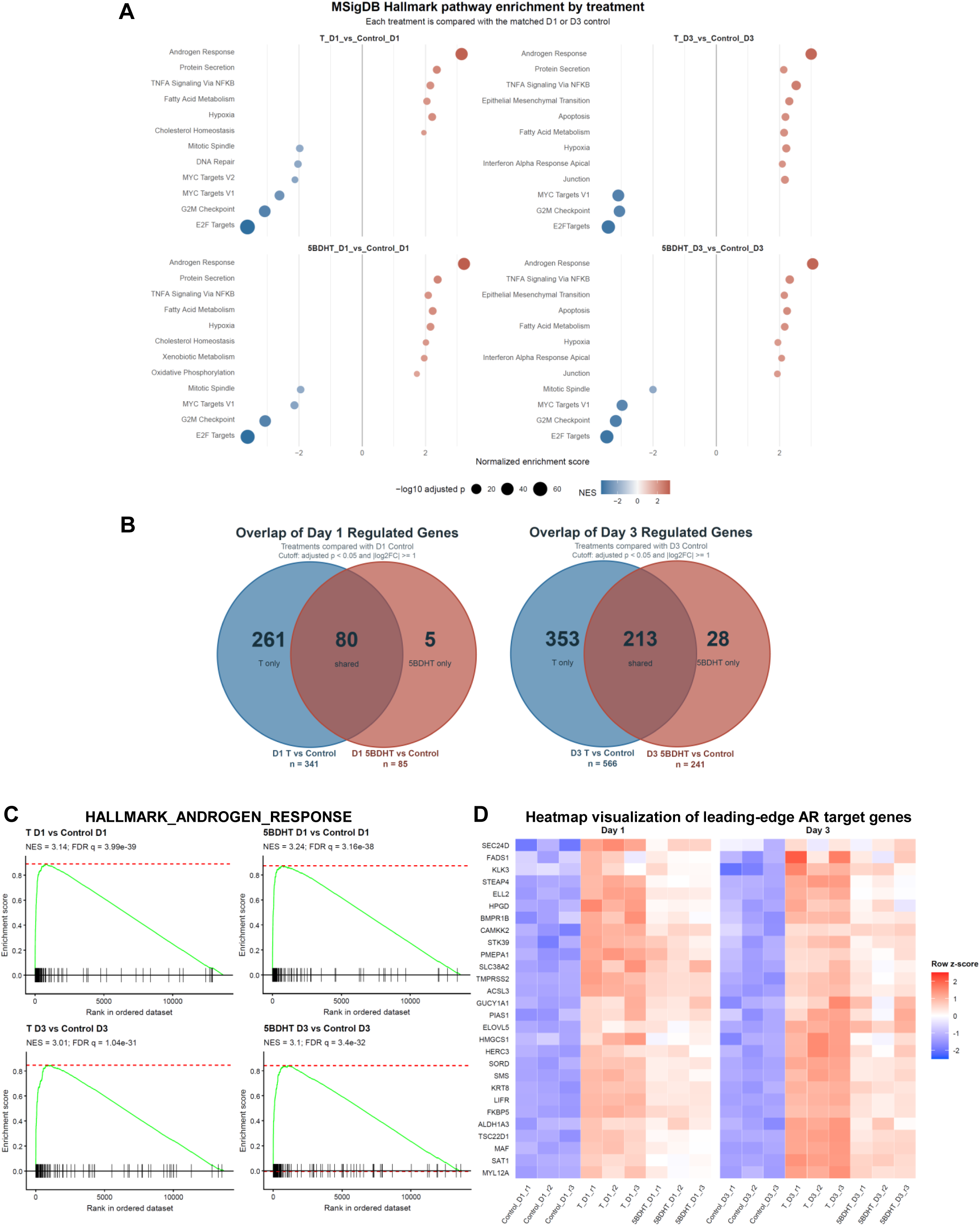
Bulk RNA-seq confirms that supraphysiologic 5β-DHT recapitulates the T androgen response transcriptome under BAT conditions. C4-2 cells in FBS medium were treated with 100 nM T or 5β-DHT and harvested at Day 1 (D1) or Day 3 (D3) (n = 3 per condition). (**A**) Hallmark GSEA dot plots at D1 and D3. (**B**) Venn diagrams of DEGs: D1 (T: 341, 5β-DHT: 85, shared: 80) and D3 (T: 566, 5β-DHT: 241, shared: 213). (**C**) GSEA enrichment plots for HALLMARK_ANDROGEN_RESPONSE at D1 and D3. (**D**) Heatmap of leading-edge AR target genes (row z-score).

Alongside AR activation, GSEA revealed significant enrichment of the cellular senescence gene set (FRIDMAN_SENESCENCE_UP [44]) by both T and 5β-DHT at D1 and D3 (T D1: NES = 1.64, FDR q = 0.005; 5β-DHT D1: NES = 1.54, FDR q = 0.033; T D3: NES = 2.25, FDR q = 3.26×10□□; 5β-DHT D3: NES = 2.1, FDR q = 5.79×10□□), with the enrichment pattern shifting from a mixed leading/trailing edge at D1 to a more pronounced leading-edge enrichment by D3, reflecting progressive deepening of the senescence transcriptional program (Fig. 5A). Heatmap analysis further showed coordinated regulation of senescence-associated genes by both ligands, including induction of CDK inhibitors and senescence-associated markers together with reduced expression of mitotic regulators such as *LMNB1* and *CCNA2*, with more pronounced changes at D3 (Fig. 5B). RT-qPCR validation confirmed these RNA-seq findings, showing significant D3 induction of *ALDH1A3*, *RRAS*, *CDKN1A*, and *CDKN2B* and downregulation of *LMNB1* and *CCNA2* by testosterone and 5β-DHT, supporting activation of a senescence-like growth-arrest program (Fig. 5C).

**Fig. 5.**
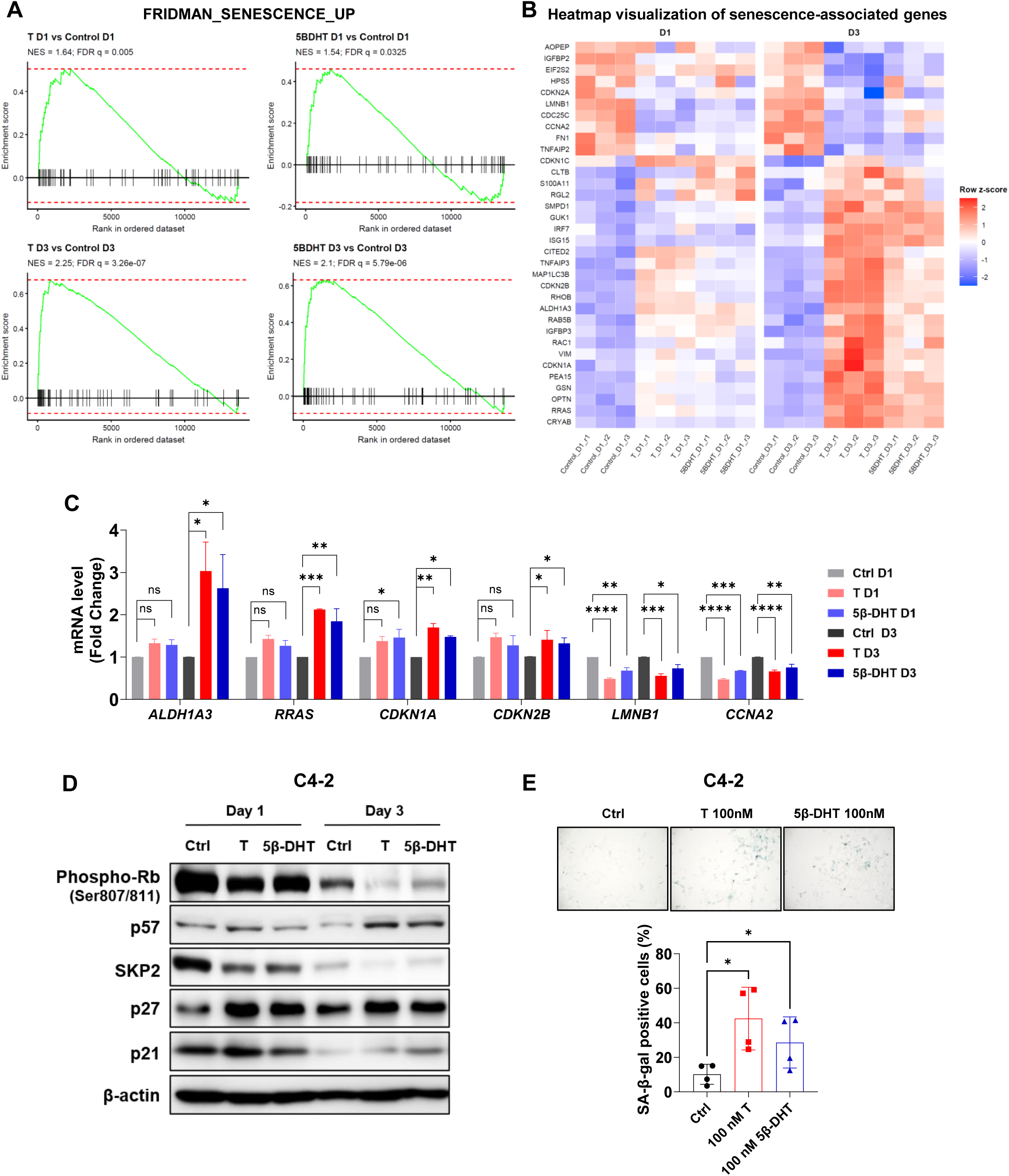
Supraphysiologic 5β-DHT induces a cellular senescence-like phenotype in C4-2 cells. C4-2 cells in FBS medium were treated with 100 nM T or 5β-DHT for D1 or D3. (**A**) GSEA enrichment plots for FRIDMAN_SENESCENCE_UP [44] at D1 and D3. (**B**) Heatmap of senescence genes (row z-score). (**C**) RT-qPCR validation for RNA-seq findings. (**D**) Western blots for phospho-Rb (Ser807/811), p57, SKP2, p27, p21, and β-actin after treatment at D1 and D3. (**E**) SA-β-gal staining in C4-2 cells treated with 100 nM T or 5β-DHT for D3 in FBS medium.

### 3.5. Supraphysiologic 5β-DHT induces senescence-associated protein changes and senescence-associated β-galactosidase (SA-β-gal) positivity

To validate the senescence transcriptional findings at the protein level, we performed western blot analysis on C4-2 cells treated with 100 nM T or 5β-DHT in FBS medium at D1 and D3. Both compounds markedly upregulated p57, p27, and p21 protein levels and concurrently reduced Rb phosphorylation at Ser807/811, indicating Rb dephosphorylation and cell cycle exit, and decreased SKP2, the F-box protein that targets p27 for proteasomal degradation. These coordinated changes are canonical markers of CDK inhibitor-driven G1 arrest and cellular senescence, and the responses to 5β-DHT closely paralleled those to T at both time points (Fig. 5D). SA-β-gal staining confirmed senescence-associated β-gal activity in T- and 5β-DHT-treated C4-2 cells at D3 (Fig. 5E). In PC-3 cells expressing AR W742C or AR H875Y, supraphysiologic 5β-DHT similarly induced the senescence protein signature, extending the mechanism to the AR mutant CRPC context (Fig. S5).

## 4. Discussion

In this study, we identify 5β-DHT as an overlooked testosterone metabolite with biologically meaningful activity in AR-mutant prostate cancer. Although 5β-DHT has historically been regarded as non-androgenic or genomically inactive due to its bent 5β-steroid conformation, our data show that it is not inert in prostate cancer models with altered AR signaling. At low androgen concentrations, 5β-DHT activated canonical AR target genes, promoted AR-dependent growth of C4-2 and LNCaP cells (both of which harbor the AR T878A mutation), and induced androgen-response transcriptional programs that substantially overlapped with testosterone. At supraphysiologic concentrations, 5β-DHT switched from supporting growth to suppressing proliferation, recapitulating key features of the BAT response. These effects extended to PC-3 cells expressing clinically relevant AR gain-of-function mutants W742C and H875Y, supporting the concept that 5β-DHT exposes a mutant-AR vulnerability in prostate cancer.

A central finding is that 5β-DHT behaves as an attenuated testosterone mimetic. In androgen-depleted conditions, both 5β-DHT and 3β-ecdiol stimulated proliferation and activated AR-regulated transcription, but their activity was weaker than that of testosterone. RNA-seq confirmed that these metabolites did not induce a distinct transcriptional program; instead, they engaged the same AR-driven regulon at reduced magnitude. This was especially clear from the differential expression analysis, where most 5β-DHT-regulated genes overlapped with testosterone-regulated genes, and all 3β-ecdiol-regulated genes were shared with testosterone. Thus, 5β-DHT and 3β-ecdiol should not be viewed as inactive terminal metabolites, but rather as weak AR agonists whose biological activity becomes evident in AR-dependent prostate cancer cells.

Importantly, the high-dose experiments revealed a functional distinction between weak AR activation and sufficient AR activation to trigger BAT-like growth suppression. Although both 5β-DHT and 3β-ecdiol promoted cell growth at low concentrations, only 5β-DHT suppressed growth in C4-2 and LNCaP cells at supraphysiologic doses. Similarly, progesterone-class steroids promoted growth at low doses but failed to suppress growth at high doses. These findings support a potency-threshold model in which AR agonist activity must exceed a minimum level to engage the growth-suppressive BAT program. This model is mechanistically consistent with the emerging view of AR as a dose-dependent transcriptional rheostat: below a critical activation level, AR drives mitogenic programs, whereas above this level, AR engages anti-proliferative and pro-senescent transcriptional programs that override growth signaling [15–17]. Our RNA-seq data directly support this interpretation, showing that supraphysiologic 5β-DHT concurrently enriches androgen-response and senescence gene sets while negatively enriching proliferative pathways, including G2M Checkpoint, MYC Targets V1, and E2F Targets, a transcriptional pattern qualitatively similar to supraphysiologic testosterone. The failure of weaker AR agonists, including 3β-ecdiol and progesterone-class steroids, to recapitulate this growth-suppressive phenotype suggests that the senescence-inducing arm of the AR transcriptional response has a higher activation threshold than the mitogenic arm, and that 5β-DHT uniquely satisfies this threshold among the testosterone metabolites and progesterone-class steroids tested here.

The activity of 5β-DHT in AR-mutant models is particularly relevant to CRPC. Ligand-binding-domain mutations can broaden AR ligand specificity and alter responses to steroids and antiandrogens [38, 39]. Consistent with this concept, 5β-DHT activated AR reporter activity in PC-3 cells expressing either AR-W742C or AR-H875Y and suppressed growth in both mutant-AR models at supraphysiologic doses. These results suggest that reduced activity at wild-type AR does not necessarily predict inactivity against mutant AR. Instead, mutant ARs may permit productive activation by ligands that are weak or poorly compatible with the canonical wild-type AR ligand-binding pocket. More broadly, these findings suggest that testosterone metabolites historically dismissed as inactive should be re-evaluated across the mutant AR repertoire found in therapy-resistant prostate cancer. This also provides a mechanistic basis for revisiting understudied steroid metabolites as potential modulators of AR signaling in CRPC.

Mechanistically, supraphysiologic 5β-DHT closely paralleled testosterone in activating AR signaling while suppressing proliferative transcriptional programs. Under BAT-like conditions, both testosterone and 5β-DHT strongly enriched the androgen-response pathway at D1 and D3, while negatively enriching the G2M checkpoint, E2F targets, and MYC-associated programs. This pattern is consistent with prior reports that supraphysiologic androgen suppresses CRPC through AR-dependent disruption of cell-cycle progression, DNA replication licensing, MYC signaling, and DNA damage responses [16–28]. The convergence between testosterone and 5β-DHT strengthened over time, suggesting that 5β-DHT can drive sustained AR-dependent reprogramming similar to testosterone, albeit with a lower overall transcriptional amplitude.

Our senescence analyses further support the idea that 5β-DHT reproduces the growth-suppressive mechanism of supraphysiologic androgen. GSEA showed significant enrichment of the FRIDMAN_SENESCENCE_UP gene set after both testosterone and 5β-DHT treatment, with stronger enrichment at D3. Heatmap and RT-qPCR analyses confirmed the coordinated regulation of senescence-associated genes, including the induction of CDK inhibitors and the downregulation of mitotic regulators such as LMNB1 and CCNA2. At the protein level, 5β-DHT increased p21, p27, and p57, reduced SKP2, and decreased Rb phosphorylation, consistent with CDK inhibitor-driven cell-cycle exit. SA-β-gal staining further confirmed a senescence-like phenotype. Together, these data indicate that supraphysiologic 5β-DHT does not simply slow proliferation nonspecifically, but activates a defined AR-associated senescence-like growth-arrest program.

These findings have potential therapeutic implications for BAT. Current BAT uses testosterone, which activates both tumor AR and wild-type AR in normal androgen-responsive tissues, contributing to systemic androgenic effects. Because 5β-DHT is a weaker activator of wild-type AR but retains activity against mutant AR and AR-dependent CRPC models, it may offer a strategy to preserve the tumor-stressing component of BAT while reducing unwanted systemic androgenicity. This possibility is especially attractive for tumors with AR gain-of-function mutations or high AR activity, where a ligand with attenuated wild-type androgenicity may still elicit sufficient AR activation to suppress tumor growth. However, this proposed therapeutic advantage remains to be tested directly.

Several limitations should be addressed in future studies. First, the current work is primarily based on cell-line models, and the pharmacokinetics, metabolism, tissue distribution, and *in vivo* antitumor activity of 5β-DHT remain unknown. Second, although the data support reduced potency relative to testosterone in AR-mutant models, a direct functional comparison of 5β-DHT activity in wild-type AR-expressing cells was not performed; future studies using wild-type AR reporter or gene-expression assays in normal androgen-responsive cell contexts will be needed to formally establish the reduced androgenicity that underpins the proposed therapeutic advantage. Third, the molecular basis for mutant-AR activation by 5β-DHT remains to be defined through ligand-binding assays, structural modeling, AR chromatin-binding studies, or coactivator recruitment assays. Finally, BAT is a dynamic treatment involving cyclic androgen peaks and troughs; future in vivo studies should determine whether cyclic 5β-DHT can reproduce testosterone-based BAT while reducing systemic androgenic effects.

In summary, our study revises the functional interpretation of 5β-DHT in prostate cancer. Rather than an inert testosterone metabolite, 5β-DHT is a weak but authentic AR agonist that activates AR-dependent growth at low concentrations and induces BAT-like growth suppression at supraphysiologic concentrations. Its ability to activate and suppress growth across multiple AR-mutant prostate cancer contexts supports 5β-DHT as a candidate lower-androgenicity testosterone surrogate for BAT and provides a rationale for further preclinical and clinical evaluation.

## Acknowledgements

This work was supported by institutional startup funds from the University of Notre Dame, awarded to Jianneng Li, and by Indiana Clinical and Translational Sciences Institute Core Pilot Funding.

## CRediT authorship contribution statement

**Samual Adams:** Conceptualization, Data curation, Formal analysis, Investigation, Methodology, Validation, Visualization, Writing – original draft, Writing – review & editing. **Lindsey Phelan:** Investigation. **Therese Lewis:** Investigation. **Josie Behm:** Investigation. **Aggie Law:** Investigation. **Xianglin Shi:** Formal analysis, Visualization. **Guangyuan (Frank) Li:** Formal analysis, Visualization. **Jianneng Li:** Conceptualization, Funding acquisition, Methodology, Project administration, Resources, Supervision, Visualization, Writing – original draft, Writing – review & editing.

## Declaration of generative AI and AI-assisted technologies in the manuscript preparation process

During the preparation of this manuscript, the authors used ChatGPT to assist with proofreading, grammar correction, and typographical editing. The authors reviewed and edited the content as needed and take full responsibility for the final content of the publication.

## Data Availability

RNA-seq data generated in this study will be deposited in the NCBI Gene Expression Omnibus (GEO) upon acceptance. All other data supporting this study are available from the corresponding author upon reasonable request.

---

**Fig. S1.**
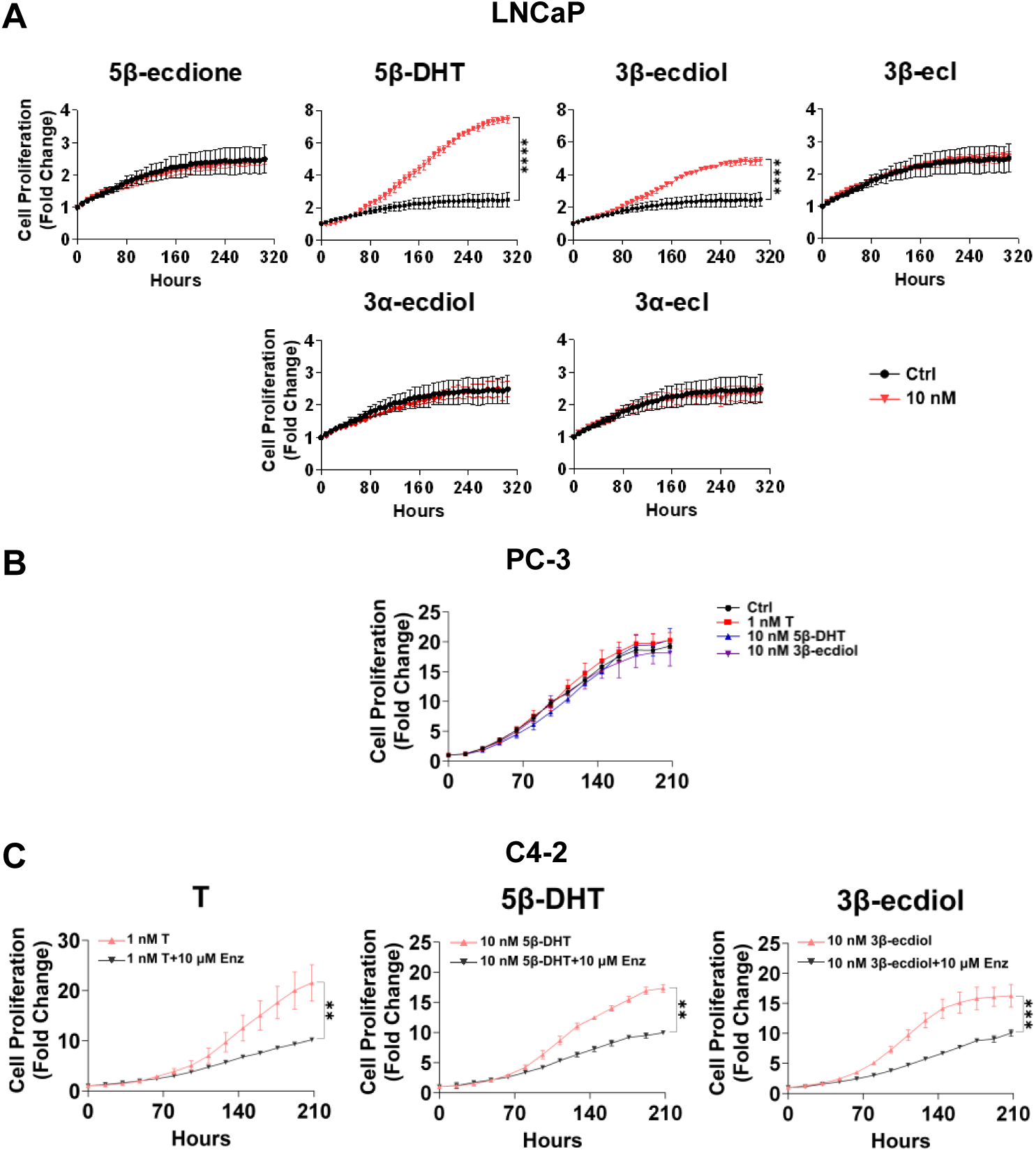
AR dependence of 5β-DHT- and 3β-ecdiol-induced cell growth and abrogation by enzalutamide. (**A**) LNCaP proliferation with 10 nM T metabolites in csFBS medium. (**B**) AR-null PC-3 parental cells show no response to T, 5β-DHT, or 3β-ecdiol. (**C**) Enz (10 μM) abrogates 5β-DHT- and 3β-ecdiol-induced growth in C4-2 cells. Data are mean ± SEM. **p < 0.01, ***p < 0.001, ****p < 0.0001.

**Fig. S2.**
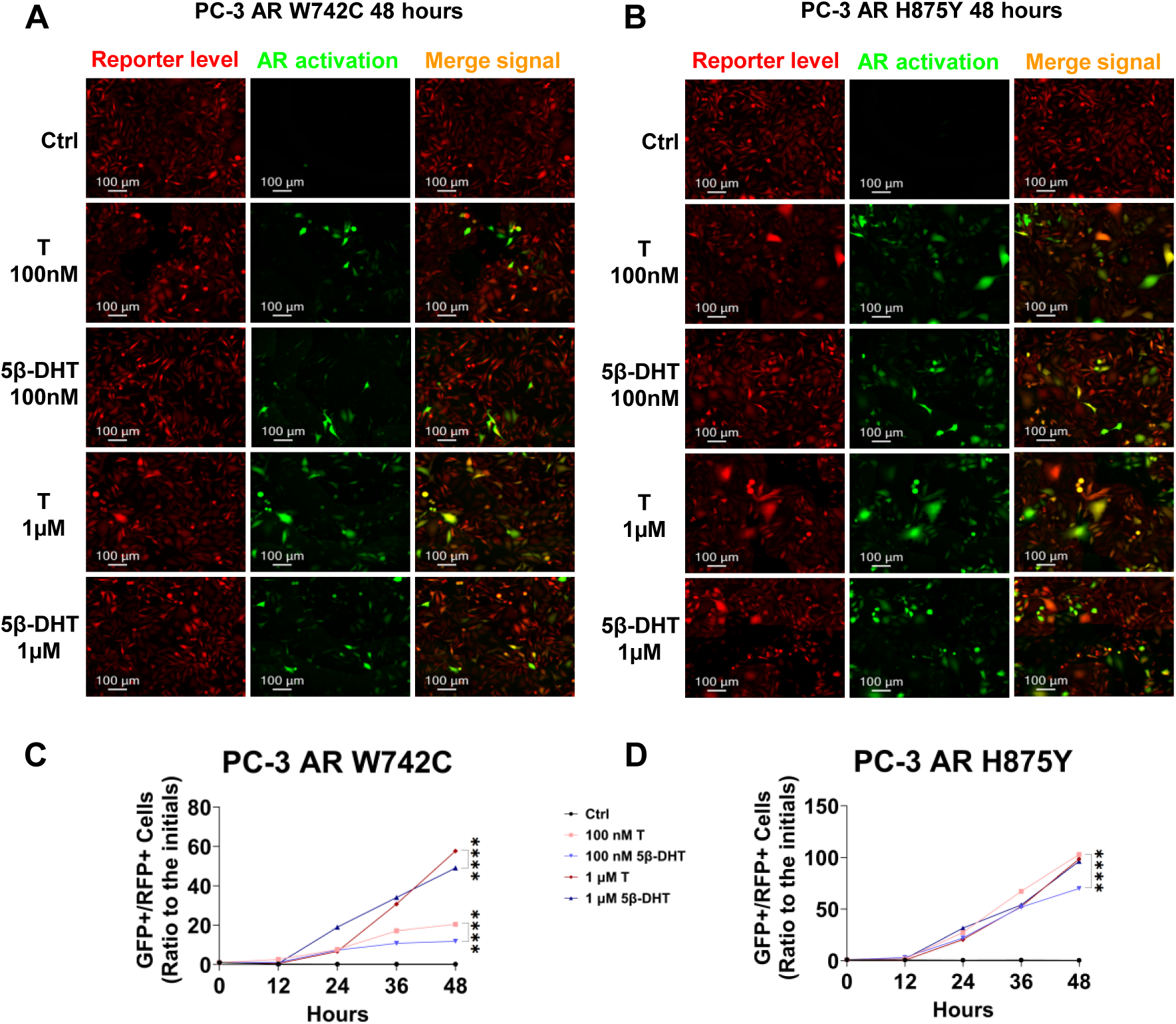
5β-DHT activates AR W742C and AR H875Y at supraphysiologic doses. PC-3 AR W742C and AR H875Y cells in FBS medium were treated with 100 nM or 1 μM T or 5β-DHT for 48 hours. (**A** and **B**) ARR3tk-eGFP/SV40-mCherry reporter fluorescence images in PC-3 AR W742C and AR H875Y cells (Ctrl, 100 nM T, 100 nM 5β-DHT). (**C** and **D**) GFP+/RFP+ ratios in PC-3 AR W742C and PC-3 AR H875Y over time with Ctrl, 100 nM or 1 μM T or 5β-DHT. Data are mean ± SEM. ****p < 0.0001.

**Fig. S3.**
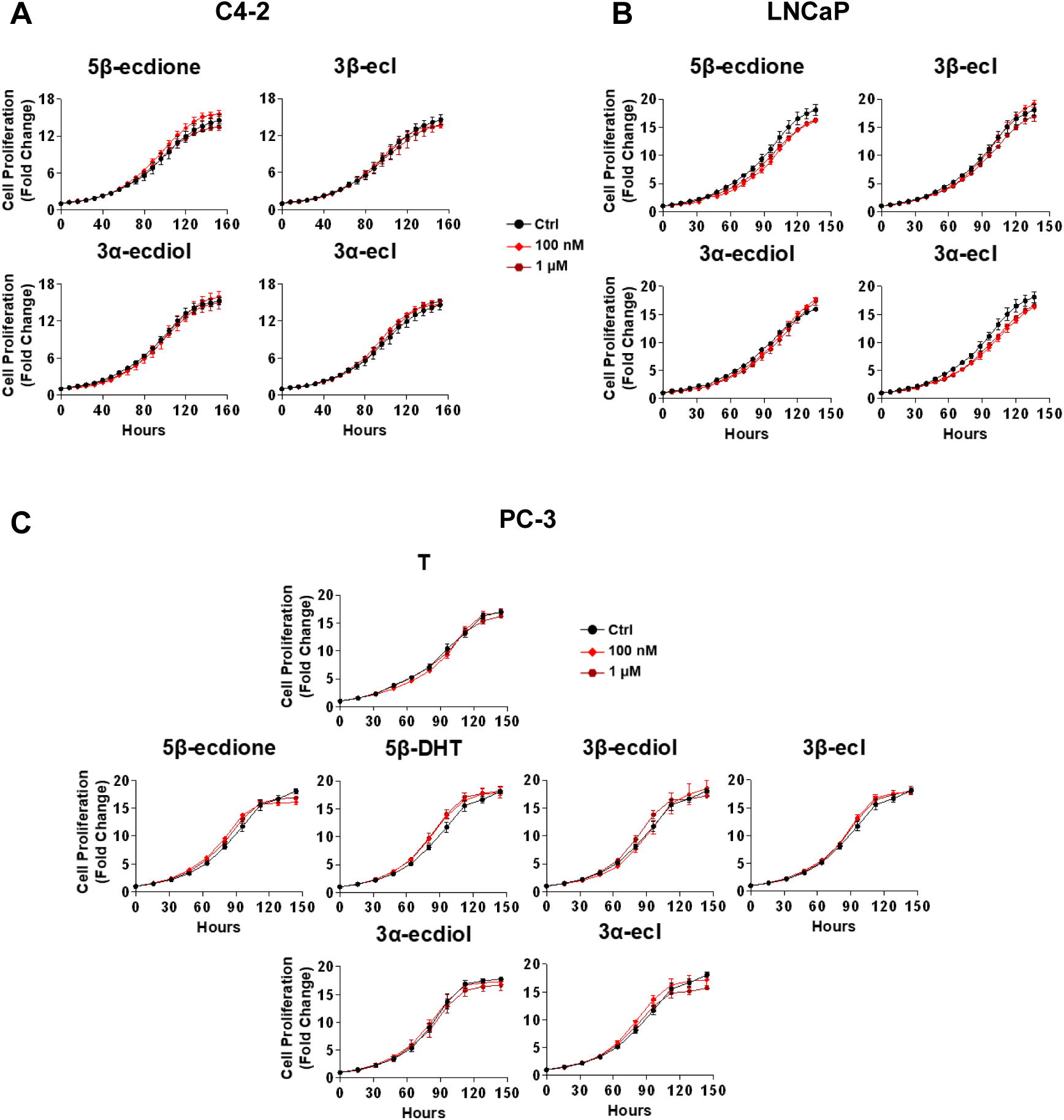
AR specificity of supraphysiologic growth suppression. (**A** and **B**) C4-2 and LNCaP cells treated with inactive metabolites at 100 nM or 1 μM in FBS show no growth suppression. (**C**) AR-null PC-3 cells are unaffected by all compounds at supraphysiologic doses.

**Fig. S4.**
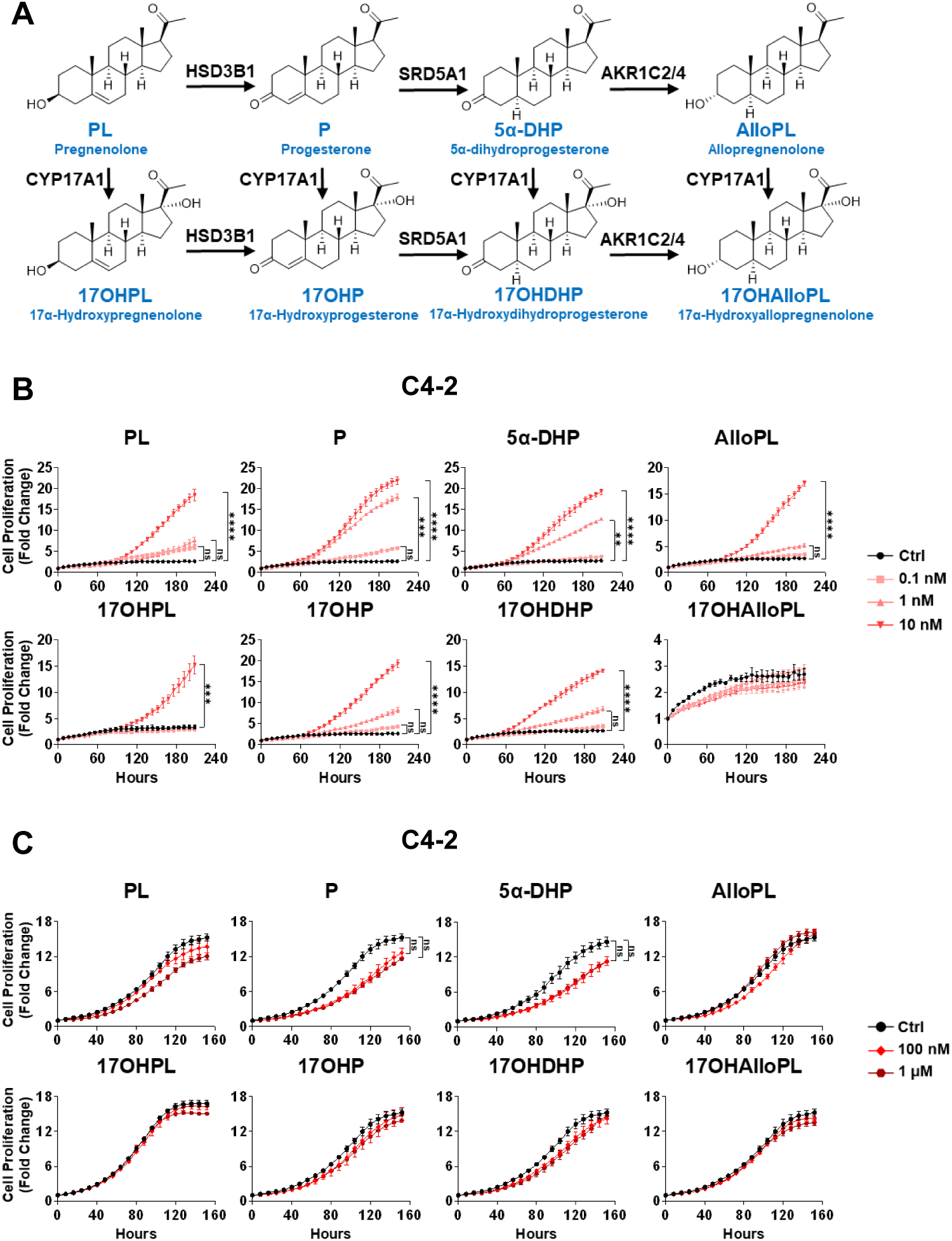
Progesterone analogs cannot suppress growth at supraphysiologic concentrations. (**A**) Progesterone metabolic pathway schematic. (**B**) C4-2 growth in csFBS medium with progestins at 0.1–10 nM. (**C**) C4-2 growth in FBS medium with progestins at 100 nM or 1 μM: none suppresses proliferation.

**Fig. S5.**
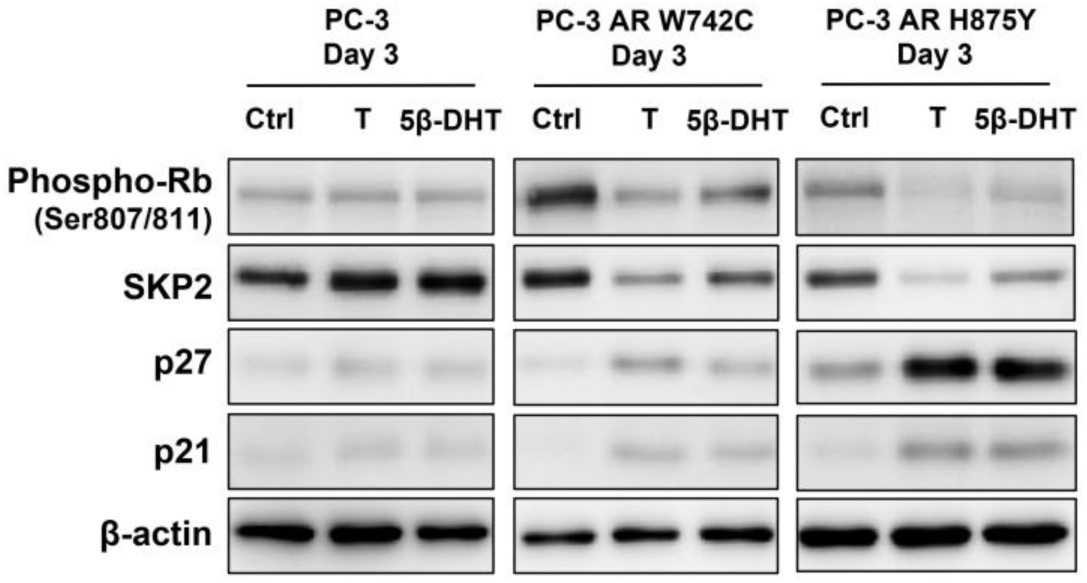
5β-DHT induces senescence-like markers in PC-3 AR mutant cells. Western blot in PC-3 parental cells, PC-3 AR W742C, and PC-3 AR H875Y cells treated with 100 nM T or 5β-DHT for D1 or D3 in FBS medium.

